# Sample pooling methods for efficient pathogen screening: Practical implications

**DOI:** 10.1101/2020.07.16.206060

**Authors:** Tara N. Furstenau, Jill H. Cocking, Crystal M. Hepp, Viacheslav Y. Fofanov

## Abstract

Due to the large number of negative tests, individually screening large populations for rare pathogens can be wasteful and expensive. Sample pooling methods improve the efficiency of large-scale pathogen screening campaigns by reducing the number of tests and reagents required to accurately categorize positive and negative individuals. Such methods rely on group testing theory which mainly focuses on minimizing the total number of tests; however, many other practical concerns and tradeoffs must be considered when choosing an appropriate method for a given set of circumstances. Here we use computational simulations to determine how several theoretical approaches compare in terms of (a) the number of tests, to minimize costs and save reagents, (b) the number of sequential steps, to reduce the time it takes to complete the assay, (c) the number of samples per pool, to avoid the limits of detection, (d) simplicity, to reduce the risk of human error, and (e) robustness, to poor estimates of the number of positive samples. We found that established methods often perform very well in one area but very poorly in others. Therefore, we introduce and validate a new method which performs fairly well across each of the above criteria making it a good general use approach.

## Introduction

For targeted surveillance of rare pathogens, screenings must be performed on a large number of individuals from the host population to obtain a representative sample. For pathogens present at low carriage rates of 1% or less, a typical detection scenario involves testing hundreds to thousands of samples before a single positive is identified. Although advances in molecular biology and genomic testing techniques have greatly lowered the cost of testing, the large number of negative results still renders any systematic pathogen surveillance program inefficient in terms of cost, reagents, and time. These costs can quickly become prohibitively expensive in resource-poor settings (e.g. pathogen surveillance in developing countries [1, 2], in non-human systems, such as wildlife disease surveillance [3]), or when reagents become scarce due to a rapid spike in testing demand (e.g. during the SARS-CoV-2 pandemic [4]).

Robert Dorfman first introduced a method to improve the efficiency of large-scale pathogen screening campaigns during World War II. In an effort to screen out syphilitic men from military service, the US was performing antigen-based blood tests on millions of specimens in order to detect just a few thousand cases. The large number of negative tests struck Dorfman as being extremely wasteful and expensive and he proposed that more information could be gained per test if many samples were pooled together and tested as a group [5]. If the test performed on the pooled samples was negative (which was very likely), then all individuals in the group could be cleared using a single test. If the pooled sample was positive, it would mean that at least one individual in the sample was positive and further testing could be performed to isolate the positive samples. This procedure had the potential to dramatically reduce the number of tests required to accurately screen a large population and it sparked an entirely new field of applied mathematics called group testing.

Due to practical concerns, Dorfman’s group testing approach was never applied to syphilis screening because the large number of negative samples had a tendency to dilute the antigen in positive samples below the level of detection [6]. Despite this, sample pooling has proven to be highly effective when using a sufficiently sensitive, often PCR-based, diagnostic assay. In fact, ad hoc pooling strategies have long been used to mitigate the costs of pathogen detection in disease surveillance programs. For example, surveillance of mosquito vector populations in the U.S. involves combining multiple mosquitoes of the same species (typically 1 – 50) into a single pool, prior to testing for the presence of viral pathogens [7–10]. Elsewhere, such pooling techniques have been successful in reducing the total number of tests in systems ranging from birds [11], to cows [12], to humans [13–15]. In many wildlife/livestock surveillance programs, sample pooling is used to simply determine a collective positive or negative status of a population (e.g. a herd or flock) without identifying individual positive samples. While this is often appropriate and sufficient for small-to-medium scale research experiments or surveillance programs, a well designed pooling scheme can easily provide this valuable information with little additional cost. For the purposes of this paper, we will focus on pooling methods that provide accurate classification of each sample so that infected individuals can be identified.

Group testing theory primarily focuses on minimizing the number of tests required to identify positive samples and many nearly-optimal strategies for sample pooling have been described. From a combinatorial perspective, a testing scheme begins by examining a sample space which includes all possible arrangements of exactly *k* positive samples in *N* total samples. Because the positive samples are indistinguishable from negative samples, a test must be performed on a sample or a group of samples in order to determine their status. The test is typically assumed to always be accurate, even when many samples are tested together (in practice, this is often not the case and approaches that consider test error and constraints on the number of samples per pool have been examined [16, 17]). In the worst case, all of the samples would need to be tested individually requiring *N* tests. The goal of group testing is to devise a strategy which tests groups of samples together in order identify the positive samples in fewer than *N* tests. Group testing methods are generally more efficient when positive samples are sparse. As the number of positive samples increases, the number of tests will eventually exceed individual testing for all of the methods. This point has been previously estimated to be roughly when the number of positives is greater than 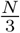 for sufficiently large *N* [18, 19]. In order to establish the most optimal testing procedure, many group testing schemes are modified based on the expected number of positive samples,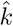. Because it is impossible to know the exact number of positive samples, problems arise when this estimate is not accurate (e.g. overestimation may require more tests to be performed than necessary, and underestimation may result in positive samples going undetected). Therefore, it is important to not only consider how different schemes scale as the number of positive samples increases but also how robust they are when the number of positive samples is misestimated.

For real-world applications, many factors should be considered when designing a pooling strategy, depending on the circumstances. Finding the best strategy often involves weighing the tradeoffs between the following factors: (a) number of tests, to minimize costs and save reagents, (b) number of sequential steps, to reduce the time it takes to complete the assay, (c) number of samples per pool, to avoid the limits of detection, (d) simplicity, to reduce the risk of human error, and (e) robustness, to poor estimates of the number of positive samples. We have identified several pooling strategies that perform well or optimally with respect to at least one of these factors. The goal of this paper is to directly compare the strengths and weaknesses of each strategy and identify the approaches that we feel are most appropriate for small-to-medium scale research experiments or surveillance programs. With this goal in mind, we favored strategies that provided the best balance across each of our criteria, particularly those that maximized the ease of performing the pooling procedure using standard laboratory equipment (i.e. defining pooling groups in ways that are easily captured using multi-channel pipettes).

We present the pros and cons of different pooling strategies by providing graphical results from computational simulations with minimal use of mathematical formulas. We focused on making the simulation results as directly comparable as possible and used realistic sample sizes (in multiples of 96 well plates) for small to medium scale experiments. The computational simulations allow us to directly compare (a) the number of tests, (b) the number of steps, (c) the number of samples per pool, (d) the number of individual pipettes, and (d) the robustness for five existing pooling strategies. We also introduce a new strategy that provides key advantages in simplicity and provides the best balance between the other criteria. Finally, we experimentally validate our strategy by testing pools of cow’s milk to detect samples that are positive for the pathogen *Coxiella burnetti*.

### Review of pooling strategies compared in this work

Pooling strategies often take either a non-adaptive or an adaptive approach. In non-adaptive methods, an optimal pooling strategy is designed in advance (for a given number of samples with an expected number of positives) and therefore it does not adapt based on information gained from the test results. Tests are run on each of the pools in parallel and the results are decoded when the tests are complete to determine which are positive. The ability to run all of the tests in parallel can save a lot of time and this is one of the main benefits of non-adaptive tests. Adaptive methods, on the other hand, require a series of steps that must be performed sequentially because each step relies on information gained from the outcome of a previous step. However, because more information is known at each step, adaptive algorithms often require fewer tests than non-adaptive methods. Below we describe several examples of both non-adaptive and adaptive pooling approaches and, in each case, we assume that the test applied to the pools is noiseless (the test will always be positive if a positive sample is present in the pool and negative otherwise) and it produces only a binary or two-state outcome (e.g. positive/negative or biallelic SNP typing).

#### DNA Sudoku

DNA-sudoku is a popular example of an optimal non-adaptive pooling strategy. This strategy is based on the idea that if a sample is present in multiple positive pools and not in any negative pools, then it is likely to be positive. However, ambiguous results can arise if multiple samples co-occur within the same positive pools because it is no longer possible to determine if one or both of the samples are truly positive. DNA Sudoku provides a more rigorous approach to avoid such ambiguity by minimizing the number of times any two samples are included in the same pool [20]. This is achieved by staggering the samples that are added to each pool in different sized windows or intervals (Fig 1); importantly, the size of the windows must be greater than 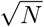 and co-prime to minimize the intersections between samples. The number of different pooling windows (the weight) should be one greater than the expected number (upper bound) of positive samples, 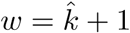, to ensure accurate results.

**Fig 1.**
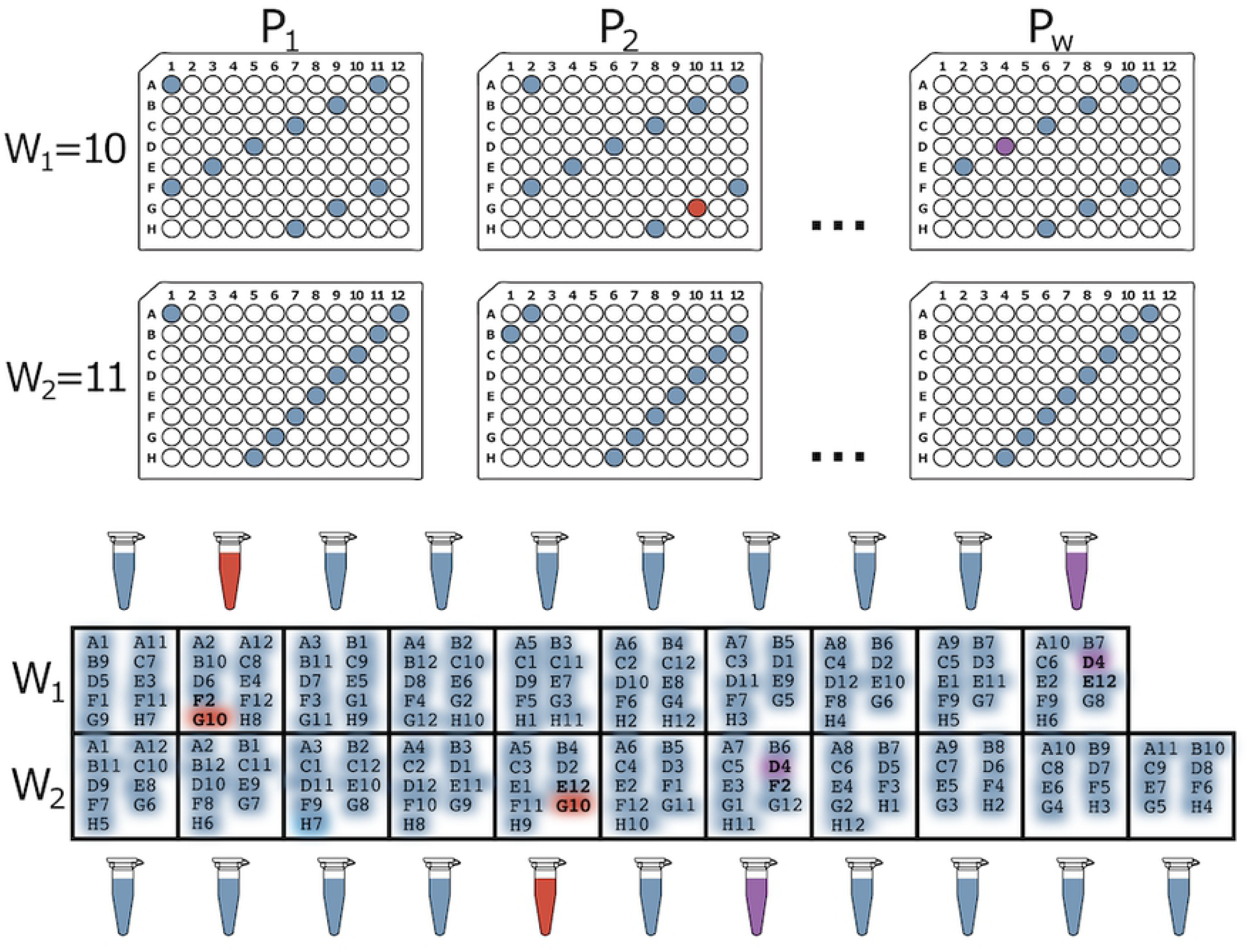
DNA Sudoku pooling example. In this example, there are a total of *N* = 96 samples. The 96-well plates show which samples are combined into each pool for the two different window sizes (*W*_1_ = 10 and *W*_2_ = 11 which are greater than 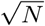 and co-prime). By using two different window sizes, the weight of this pooling design is *w* = 2 meaning that *k* = *w* − 1 = 1 positive sample can be unambiguously identified in a single step using *T* = *W*_1_ + *W*_2_ = 21 tests. The positive samples are decoded by finding the samples that appear most often in the positive pools. For example, if G10 is the only positive sample, we can detect this from the pooling results by noticing that G10 was added to both of the positive (red) pools while other samples in those pools were added to only one or the other. Alternatively, if both G10 and D4 are positive, four samples occur with equal frequency (D4, G10, E12, and F2) in the positive pools (red and purple) and it is impossible to determine which are the true positive samples. This ambiguity is introduced because the test was designed to handle only one positive sample.

Once the samples are pooled for each window size and the pools are tested, a decoding scheme is used to identify the positive samples from the positive pools. The decoding works by identifying which samples occur the most frequently in the positive pools. If the weight is chosen correctly using a good estimate of *k*, all of the positive samples can be unambiguously identified. However, if the true number of positive samples exceeds 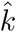, the results become ambiguous and false positives can occur. Over estimating the maximum number of positive samples can provide a buffer against ambiguous results but this comes with a large increase in the number of tests (*> N* additional tests for each additional pooling window). Alternatively, the ambiguous samples can be tested individually, but this requires an additional round of testing which voids one of the main advantages of non-adaptive testing.

When 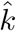 is estimated appropriately, DNA Sudoku is a very efficient non-adaptive approach, especially when the number of samples is very large. It was originally designed for pooling and barcoding thousands of DNA samples in preparation for high-throughput sequencing. However, because it was intended for use in large-scale sequencing facilities with robotic equipment, the pooling design is complex and intricate and therefore difficult for a human technician to perform accurately and consistently by hand.

#### Two Dimensional Pooling

Multidimensional pooling is another non-adaptive approach that is generally easier to perform than DNA Sudoku but can be more prone to producing ambiguous results. As the name implies, this procedure can be extended to many dimensions [21, 22], however it becomes more difficult to perform without robotics when more than two dimensions are used. In the two dimensional (2D) case, *N* samples are arranged in a perfectly square 2D grid or in several smaller but still square sub-grids [23]. For example, when testing 96 samples (as in Fig 2), this could be achieved through a single 10×10 grid or through 4 5×5 sub-grids (with 4 empty spaces). Once arranged, all of the samples along each individual column and each individual row are pooled. This results in 20 pools for a 10×10 grid, and 40 pools for 5×5×4 grids. Once the pools are tested, the positive samples are decoded by identifying which of them are present at the intersection of positive rows and columns [23].

**Fig 2.**
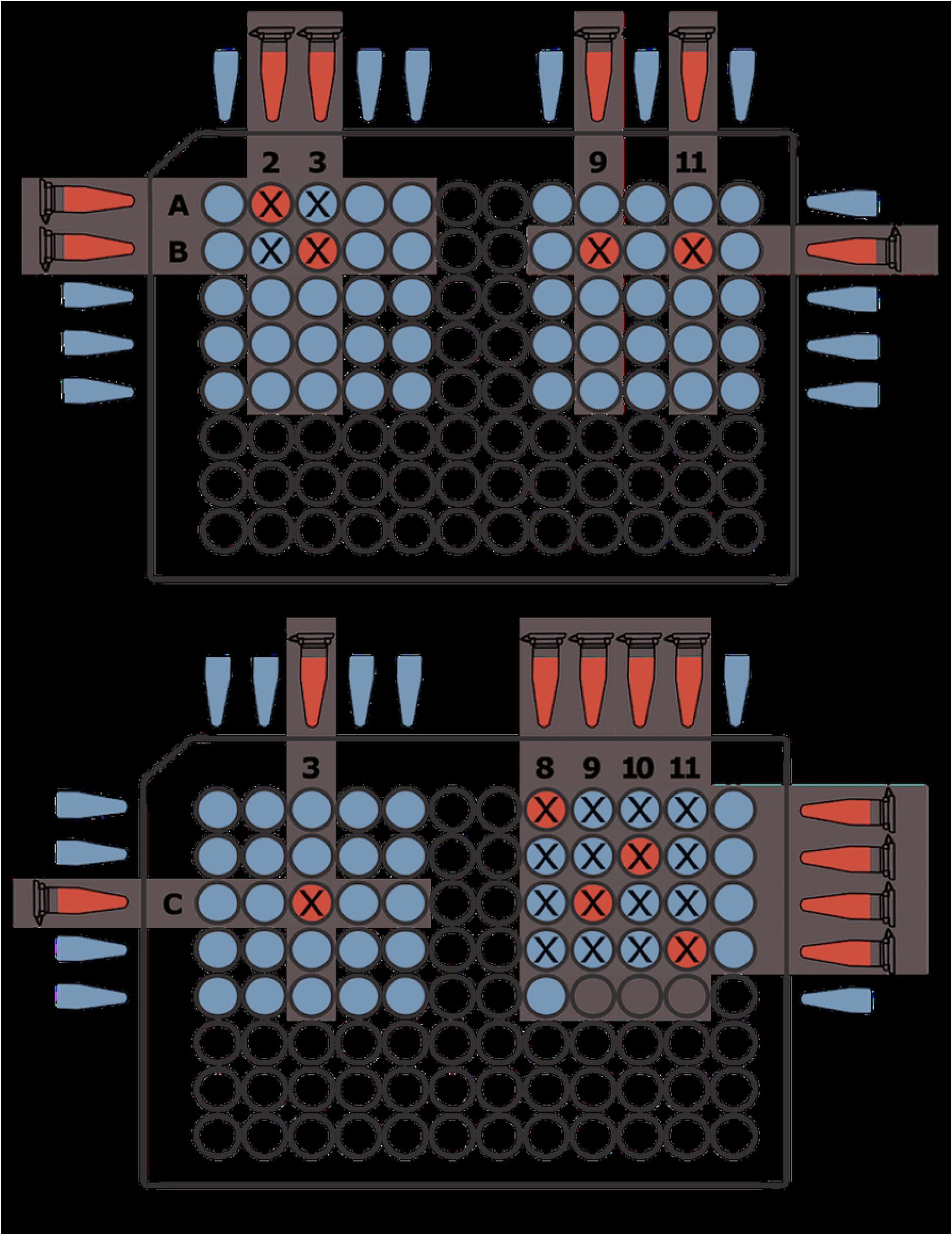
Two-Dimensional pooling example. A total 96 samples are arrayed in symmetrical 5×5 grids (with 4 empty wells in the last grid) and *k* = 9 of the samples are positive (red wells). The pooling procedure combines each row and each column of a grid into separate pools for a total of *T* = 2 × 5 × 4 = 40 tests. Samples that are at the intersection of a positive row and a positive column (marked with an “X”) are potentially positive samples. When more than one row *and* more than one column are positive, some of the samples at the intersections are likely false positives (e.g. the top left and bottom right grids). Otherwise, the results are unambiguous and the correct positive samples can be identified (e.g. the top right and bottom left grids).

In 2D pooling, ambiguous results arise when positive samples are present in multiple rows *and* multiple columns (e.g. in the top left grid in Fig 2, the two positive rows and the two positive columns intersect at four wells, only two of which (red wells) are positive). When this occurs, the number of intersecting points is almost always higher than the true number of positive samples. Ambiguous results can be somewhat mitigated by decreasing the size of the grid as the expected number of positive samples increases. The chances of ambiguous intersections increase when there are more positive samples, so using more grids of smaller size (and consequently more tests), will make ambiguous arrangements less likely. Alternatively, the ambiguous samples can be tested individually in a second followup round of testing, but this again nullifies the main benefit of non-adaptive testing, which is the ability for all tests to be carried out in parallel.

#### S-Stage Approach

Dorfman’s original pooling design for syphilis screening was an adaptive two-stage test. Following this method, samples are partitioned and tested in *g* groups of size *n*. All of the samples in groups with negative results are considered to be negative and all of the samples in groups with positive results are tested individually. Ignoring the constraints of the actual assay, the optimal group size that minimizes the number of tests depends on the number of positive samples, *k*. Specifically, there should be roughly 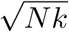 groups of size 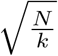 [5, 24]. Dorfman’s two-stage approach was later generalized to any number of stages using Li’s S-Stage algorithm [24], which can reduce the number of tests required to identify positive samples. At each stage, *s*_*i*_, of the S-Stage algorithm (Fig 3), the untested samples are arbitrarily divided into *g*_*i*_ groups of size *n*_*i*_ and the test is performed on each group. The samples in pools with negative test results are deemed negative and removed from consideration. The samples in positive pools move on to the next stage where they are redivided into *g*_*i*+1_ groups of size *n*_*i*+1_. This is repeated until the final stage, where *n*_*s*_ = 1, and all of the remaining samples are tested individually. The optimal number of samples per group at each step is 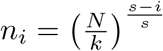 and the optimal number of steps is 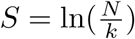 which achieves an upper bound of 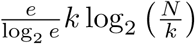 tests. Li demonstrated that misestimation of the number of positive samples, 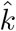, has only a small impact on the total number of tests, especially when the number of stages is high. The S-Stage algorithm can require many more steps than non-adaptive algorithms, but when the number of steps is low, it compares favorably, especially in cases when the non-adaptive methods require additional validation steps.

**Fig 3.**
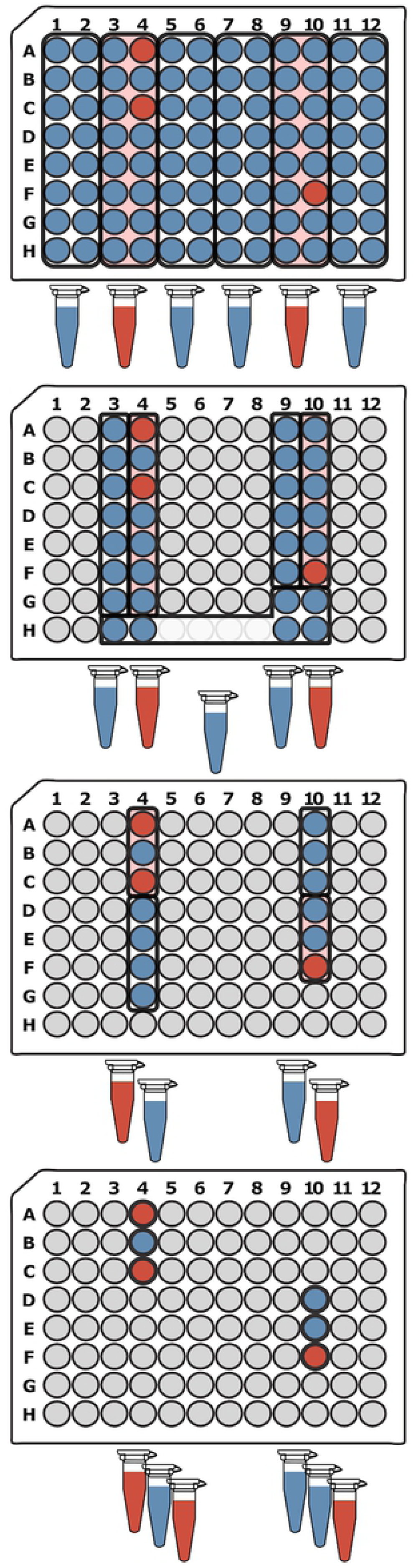
S-Stage pooling example. For 96 samples with an estimate of 3 positive samples, the S-Stage algorithm requires 4 steps. In the first step (top 96 well plate), 96 samples are tested in 6 groups (black outline) of 16. In the next step, the samples in the positive pools from the previous step are arbitrarily redivided into 5 groups of 6 or 7 samples and tested. In the third step, the samples from positive pools from step 2 are redivided into 4 groups of 3 or 4. In the final step, individual testing is performed on samples from the positive pools in step 3. The number of tests required depends on the initial arrangement of positive samples within the pools but in this example 21 tests are required to identify 3 positive samples (red wells). The number of tests is lower than the upper bound in this case due to the fortunate placement of two positive samples in the same pool in steps 1-3.

#### Binary Splitting by Halving

Sobel and Groll [25, 26], introduced several adaptive group testing algorithms based on recursively splitting samples into groups and maximizing the information from each test result. They demonstrated that this class of algorithm is robust to inaccurate estimates of *k*, particularly in the case of the Binary Splitting by Halving algorithm which can be performed without any knowledge of the number of positive samples. Binary Splitting by Halving (Fig 4) begins by testing all of the samples in a single pool. If the test is negative, all of the samples are negative and testing is complete, if the test is positive, the samples are split into two roughly equal groups and only one of the groups is tested in each step. If the tested half is negative, we know that all of the samples in the tested group are negative and testing is now complete for those samples. We also know that the untested half must contain at least one positive sample (because the test containing all of the samples tested positive). Alternatively, if the tested pool is positive, we know that it contains at least one positive sample and we know nothing about the untested half. In either case, the binary splitting always continues with the group that is known to contain a positive sample until a single positive sample is identified with individual testing. At this point, all of the samples that remain untested are added to a single pool and tested, beginning the process again. This is repeated *k* times and stops when the initial test of all the remaining samples is negative or when all samples have been tested (either through individual testing or elimination). Using this method, *k* positive samples can be identified in at most *k* log_2_ *N* tests. Binary splitting is only efficient when fewer than 10% of samples are positive, otherwise more tests are required than individual testing [25, 26]. This is the only approach discussed here that does not rely on an estimate of *k* and therefore the performance is not impacted by misestimation of the number of positive samples.

**Fig 4.**
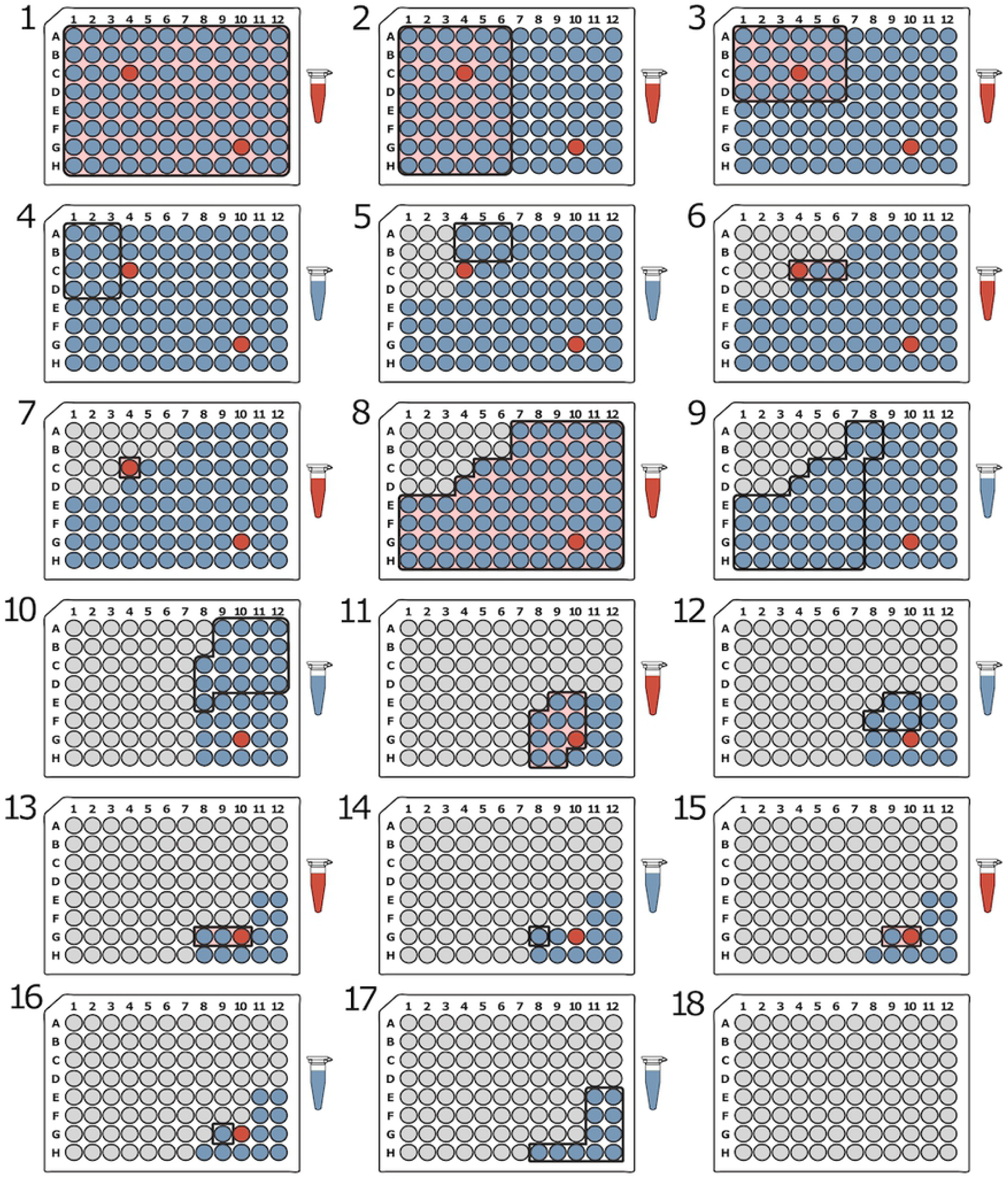
Binary splitting by halving pooling example. In this example, there are *N* = 96 samples and two of the samples are positive (red wells). To begin, all of the samples are pooled and tested (Step 1). If the first test is negative, testing is complete and all samples are considered negative. Otherwise, half of the samples are pooled and tested (Step 2). If the tested half is negative, then all of the samples in the tested half are considered to be negative and at least one negative sample is known to be present in the other non-tested half of the samples. If the tested half is positive, then it contains at least one positive sample and no information is gained about the other untested half. In either case, the method continues by halving and testing whichever group is known to contain a positive sample until a single positive sample is identified (either by individual testing, as seen in Step 7, or by elimination, as seen in Step 16). Once a single positive sample is identified, the remaining unresolved samples (non-grey wells) are pooled and tested to determine if any positive samples remain and the process continues until all positive samples are identified. Only one test is required per round, and in this example, it takes 17 sequential rounds to recover both positive samples.

#### Generalized Binary Splitting

Hwang’s Generalized Binary Splitting algorithm is very similar to Binary Splitting by Halving (Fig 4) except the size of the first split is optimized for the expected number of positive samples. This is important because it helps bypass some of the early and least productive tests. In the Halving method, as the number of positive samples increases, the first few tests are more likely to be positive due to chance. Positive tests, in general, provide the fewest pieces of information and do not eliminate any negative samples; consequently, positive tests are particularly inefficient early on when the potential to eliminate large groups of samples is highest. Additionally, it means that each binary search will begin with a large number of samples which will require more tests and steps to identify the first positive sample. To solve this problem, The Generalized Binary Splitting algorithm attempts to modify the size of the initial pool so that it is small enough to capture a single positive sample on average. When smaller groups are tested they are less likely to be overwhelmingly positive which means more samples can be eliminated in negative tests and a single positive sample can be found quicker using fewer tests [27]. As the ratio of samples to positive samples 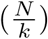 increases, the number of tests required to identify *k* positive samples approaches *k*log_2_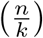 which is nearly optimal; however, like Binary Splitting by Halving, the Generalized Binary Splitting approach requires many sequential steps to complete testing.

#### Modified 3-Stage Approach

Here we are introducing a new approach that we developed with the goal of finding a good balance between the number of tests, the number of steps, simplicity, and robustness. We found that many of the methods described previously focus on optimizing only one of these features usually to the detriment of the others. Instead of attempting to perform the best in a single area, we wanted to take a more balanced approach and find tradeoffs that allow good performance across each of these areas. Our Modified 3-Stage approach (Fig 5) is based on the S-Stage approach but it is modified so that the number of steps is constrained to a maximum of three. At three steps, this approach requires only one additional step than ambiguous non-adaptive approaches that require two steps for complete validation. Because the S-Stage algorithm is already fairly robust, constraining the number of steps does not have a large impact on the number of tests required. We also modified our method to be simpler and easier to perform by borrowing the recursive subdividing used in the binary splitting approaches. In the S-Stage approach, the remaining samples in each step are arbitrarily redivided into pools. Not only does this make it difficult to keep track of the remaining samples spread across the plate, it can also make it more difficult to collect the samples for a pool using a multichannel pipette (e.g. Step 2 in Fig 3). Instead, we opted to recursively subdivide the samples from positive pools. This makes it easier to keep track of the samples that should be pooled at each stage and, because the samples are always in close proximity, they are easier to collect using a multichannel pipette (compare 3 and 5).

**Fig 5.**
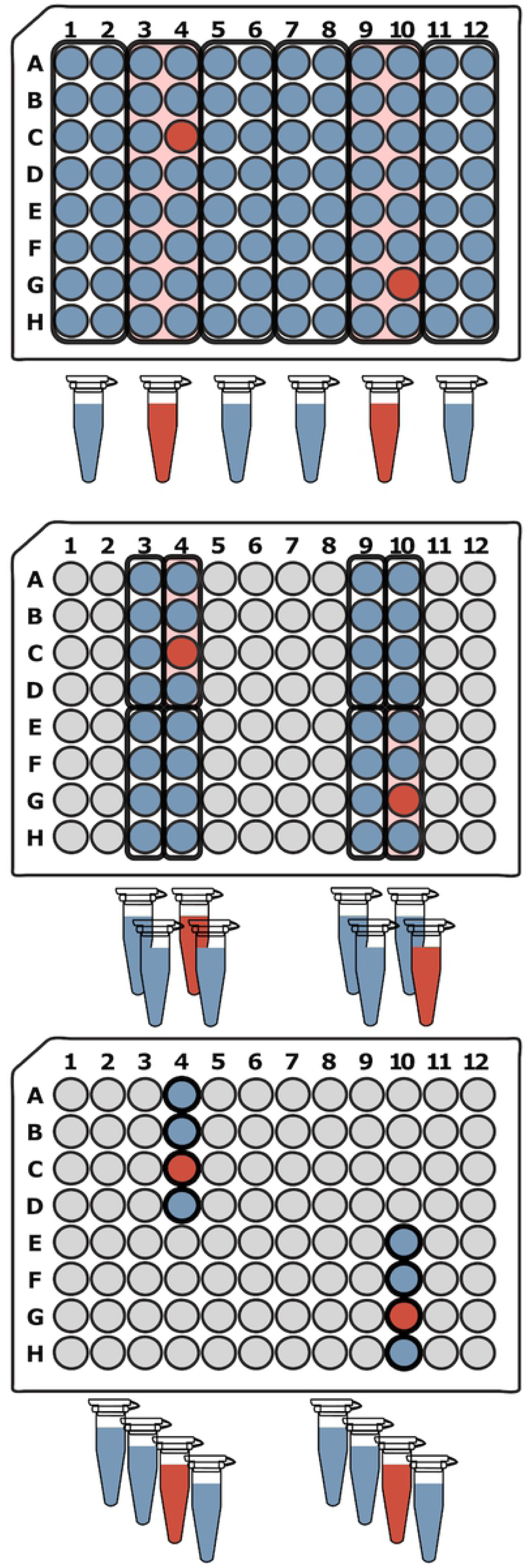
Modified 3-Stage pooling example. For 96 samples and an estimate of 2 positive samples, the Modified 3-Stage approach begins by creating 6 pools with 16 samples each. The positive pools from the first step are then subdivided into 4 groups of 4 in the second step. In the final step, the samples from the positive pools in step 2 are tested individually. In the modified 3-Stage approach, the pools are recursively subdivided into groups instead of arbitrarily redividing the remaining samples at each step. This is simpler and keeps the samples for each subsequent pool in close proximity. The total number of tests depends on the arrangement of the positive samples, but in this example, the modified 3-stage algorithm requires 22 tests.

## Materials and methods

### Computational simulations

Computational simulations were carried out for each of the six pooling strategies described above. The number of samples, *N*, in each test were in multiples of 96 to represent 96-well plates: 1 × 96 = 96, 4 × 96 = 384, and 16 × 96 = 1, 536. Each set of samples was represented as a binary array of size *N*, where 1’s represented positive samples and 0’s represented negative samples. For each test, 100 simulations were generated by placing *k* positive values in random positions in the array, with *k* ranging from 1 to 20. In each simulation, the number of tests, the number of sequential steps, and the number of individual pipettings required to make the pools were recorded. In cases where it was appropriate, the number of pipettings was calculated assuming either an 8- or a 16-channel pipette in addition to a single channel pipette. We only considered pooling schemes that were able to completely and accurately identify all of the positive samples in the sample set. To accomplish this, some of the pooling schemes required additional steps and tests that are accounted for in the simulation. The simulation code is available at https://github.com/FofanovLab/sample_pooling_sims.

#### DNA Sudoku Simulations

For the DNA Sudoku experiments, we tested different weights ranging from 2 to the highest value that did not exceed the number of tests required for individual testing. For example, with a sample size of 96, the maximum weight we used was 6 with window sizes of 10, 11, 13, 17, 19, and 23; this testing design required 93 tests, in the unambiguous case, and including any additional testing windows would cause the number of tests to exceed individual testing. The window sizes at the maximum weight were 20, 21, 23, 29, 31, 37, 41, 43, 47, and 53 for 384 samples; and 40, 41, 43, 47, 49, 51, 53, 59, 61, 67, 71, 73, 79, 83, 89, 97, 101, 103, 107, 109, and 113 for 1,536 samples. For smaller weights, the window sizes were just the first *w* window sizes listed here for each sample size. For the first round of testing, the total number of tests was equal to the sum of the window sizes.

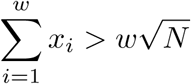

If the result was ambiguous (i.e. any time the number of positives exceeded *w* − 1), the number of steps increased to two and additional tests, equal to the number of prospective positive samples, were added to the test count. Because the samples that were added to each pool are staggered, multichannel pipettes do not provide any advantage; therefore, the number of pipettings was calculated assuming only a single channel pipette (*N* × *w* + |Prospective Positives|).

#### 2D Pooling Simulations

For the 2D pooling simulations we used square *D* × *D* grids and each of the *M* grids in a simulation were the same size. The samples were pooled along each row and column requiring 2*DM* tests. When the results were ambiguous, the number of steps increased by one and the number of tests increased by the number of prospective positive samples to account for the validation. Because the pooling along columns and rows can be easily and more efficiently performed using multichannel pipettes, the number of pipettings was calculated using an 8- and a 16-channel pipette, in addition to a single channel pipette. The number of pipettings was the 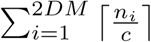 where *n*_*i*_ is the number of samples in each row or column and *c* is the number of channels in the pipette. We assumed that any additional pipettings required for testing the ambiguous samples was performed with a single channel pipette.

#### S-Stage Simulations

The S-Stage simulations were provided with an expected number of positive samples, 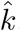. The number of steps was calculated as 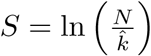 and the number of samples per group was 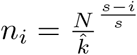. Because these calculations do not provide integer values, a nearest integer approximation was used. Optimal integer approximations of these values can be determined numerically but here we consistently applied a ceiling function. For each number of true positive samples (*k* = 1 − 20) we ran simulations with expected values, 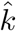, ranging from 1-20. The number of tests was calculated as

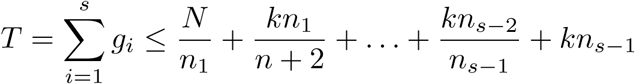

where 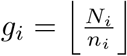 is the number of groups tested at each step. The number of pipettings for a single channel pipette was equal to the number of samples in each of the pools that were tested. For multichannel pipettes, the number of samples in each pool was divided by the number of channels and rounded up. In cases where the samples in the pool were not in adjacent wells, additional pipettings were required.

#### Modified 3-Stage Simulations

Our modified 3-Stage approach is similar to the S-Stage algorithm except that the number of steps was constrained to a maximum of three: 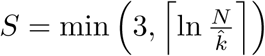. In order to recursively subdivide each pool, the number of subgroups was calculated as 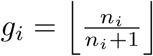 with *n*_*i*_ calculated the same way as the S-Stage simulations. For each true number of positive samples (*k* = 1 − 20) we ran simulations with expected values ranging from 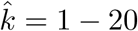. The number of tests and the number of pipettings were calculated the same way as the S-Stage simulations.

#### Binary Splitting by Halving

The Binary Splitting by Halving simulations did not require any estimate of the number of positive samples. The simulation performed repeated binary searches for positive samples until no more positive samples remained. Only one test was performed at each step and, because each step depended on information gained in the previous step, none of the steps were performed in parallel. Therefore, the number of tests was equal to the number of steps. The number of pipettings was equal to the size of each pool divided by the number of channels in the pipette, rounded up.

#### Generalized Binary Splitting

The Generalized Binary Splitting simulations were similar to the Binary Splitting by Halving simulations except that the initial group size was calculated based on the number of expected positive samples 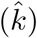. More specifically, the initial group size was calculated as 2^*a*^ where 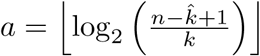. Binary Splitting by Halving (as described above) was performed on the initial group until a positive sample was identified at which point the value *N* was updated to reflect the number of remaining untested samples and the value 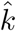 was decremented by 1 if a positive sample was found. The next group of 2^*a*^ was calculated using updated values of *N* and 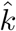. This continued until either 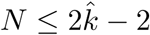, at which point the remaining samples were tested individually, or 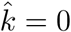, at which point all of the suspected positive samples were identified. Because the standard algorithm only guarantees that up to 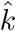 positive samples will be found, we added additional rounds of binary splitting to ensure all of the positive samples were identified. The number of tests, steps, and pipettings were calculated the same way as the Binary Splitting by Halving simulations.

### Experimental validation of modified 3-stage approach

We set up rare pathogen detection experiments in complex microbiome backgrounds to test our Modified 3-Stage approach. We used a total of 768 samples (eight 96-well plates) that contained a background of 2 *µ*L of DNA extraction from cow’s milk and 8 *µ*L of molecular grade water. These samples originated from 24 distinct cow milk samples and were replicated (32 replicates each) to fill eight 96-well plates – a total of 24 unique microbiome backgrounds. *C. burnetti* DNA (1 *µ*L) was added to 10 randomly chosen background samples (∼1.3% carriage rate) as we verified that the spike-in was successful using a highly sensitive Taqman assay designed to target the IS1111 repetitive element in *Coxiella burnetti* [28]. Using the same Taqman assay, we also verified that the target pathogen was not present in any of the 24 unique microbiome backgrounds prior to the spike-in. To ensure a consistent amount of background DNA, the milk extractions were tested to determine the amount of bacteria with a real-time PCR assay that detects the 16S gene and compares it to a known standard [29].

The pooling procedure was carried out by a typical researcher looking to identify samples that are positive for the pathogen of interest, *C. burnetti*. Assuming *N* = 96 and 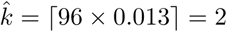 the pooling scheme recommended by our modified 3-stage approach is depicted in Fig 5. In the first step, 6 pools consisted of 16 samples each, collected along every 2 columns of the 96 well plate using an 8-channel pipette. The 2 *µ*L aliquots from each sample were collected in a plastic reservoir and then pipetted back into a single well in a new 96 well plate. The *C. burnetti* Taqman assay was used to test each of the pools. For the reaction, the following were combined for a final volume of 10 *µ*L: 1 *µ*L from the pool, 2 *µ*L of Life Technologies TaqMan® Universal PCR Master Mix for a final concentration of 1X, 0.3 *µ*L each of the forward and reverse primers for a concentration of 0.6 µM, 0.13 *µ*L of the probe for a concentration of 0.25 µm and molecular grade water to a final volume of 10 *µ*L. The reaction was run on an Applied Biosystems 7900 Real Time PCR system with the following conditions: 50 °C for 2 minutes, 95 °C for 10 minutes, and 40 cycles of 95°C for 15 seconds and 60°C for 1 minute. The second pooling step was carried out by subdividing the samples from the positive pools in the previous step into four groups of four samples. Again, 2 *µ*L from each well was combined into the pool. These pools were subjected to Taqman *C. burnetti* assay as described above. Finally, the individual samples belonging to pools positive in the second pooling step, were tested as described above.

## Results and Discussion

### Number of tests

Because minimizing the number of tests is one of the primary goals of group testing, we begin by comparing the number of tests required for each method using a range of sample sizes: 96, 384, and 1,536. The number of positive samples ranged from 1 to 20 which resulted in minimum positive rates of 1.04%, 0.26%, and 0.07%; and maximum positive rates of 20.83%, 5.20%, and 1.30% for 96, 284, and 1,536 samples, respectively. Fig 6 directly compares the average number of tests for each method using the optimal parameter settings. For the S-Stage, Modified 3-Stage, and General Binary Splitting approaches, the results shown are for simulations where the expected number of positive samples was the same as the true number of positives 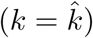. For DNA Sudoku and 2D Pooling, the results shown are for simulations with parameters that resulted in the lowest average number of tests.

**Fig 6.**
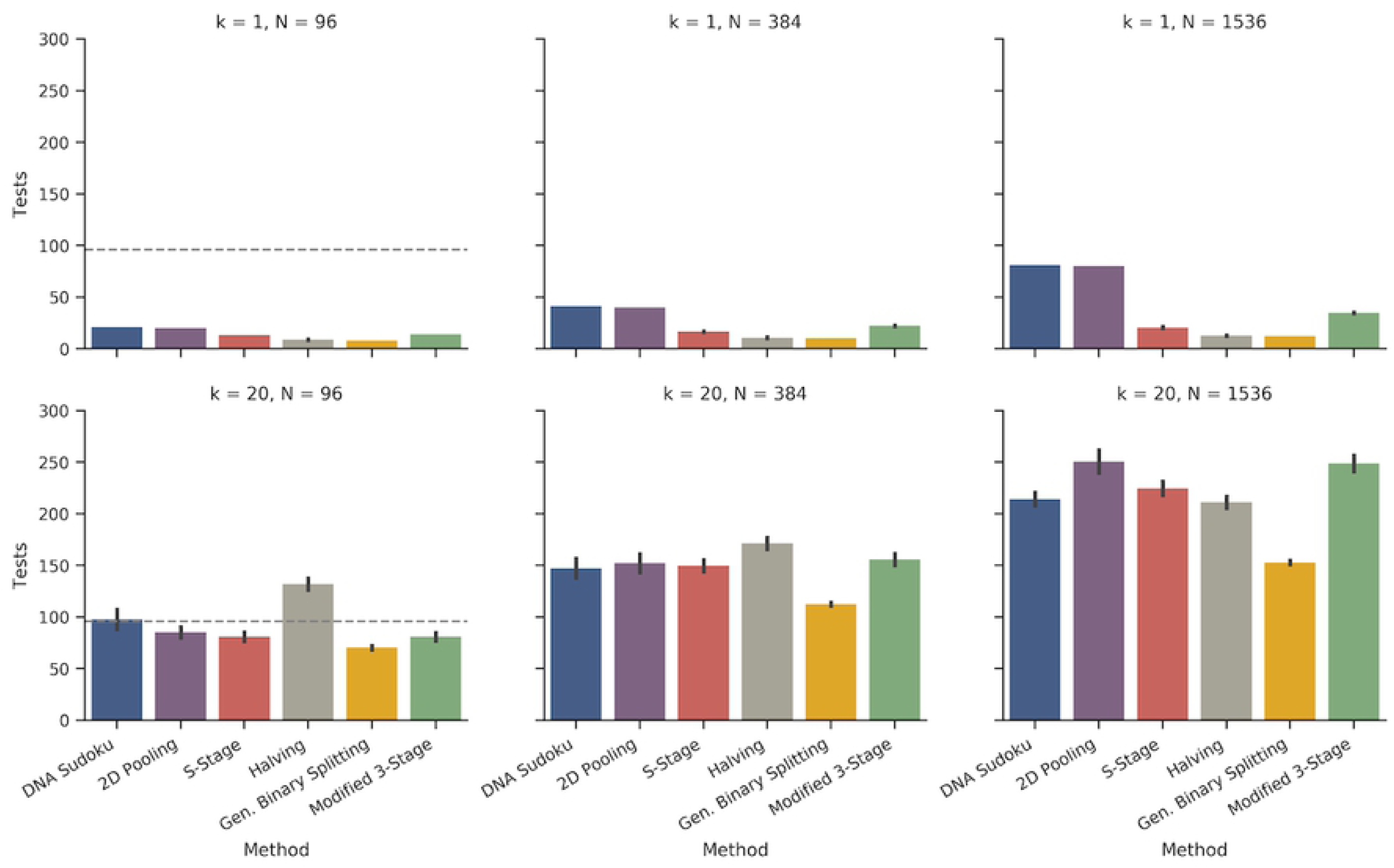
Comparison of the number of tests required for each pooling method. The bar graphs show the average number of tests required for each method and the error bars are the standard deviation across 100 simulations. The top row shows simulations with one positive sample and the bottom row shows simulations with 20 positive samples. The columns are different sample sizes from left to right: 96, 384, and 1,596. For the S-Stage, Modified 3-Stage, and General Binary Splitting approaches, the results shown are for simulations where the expected number of positive samples was the same as the true number of positives. For DNA Sudoku and 2D Pooling, the results shown are for simulations with parameters that resulted in the lowest average number of tests (DNA Sudoku: *w* = 2 when *k* = 1, and when *k* = 20, *w* = 3 for 96 samples and *w* = 4 for 384 and 1,536 samples; 2D Pooling: when *k* = 1, the grid sizes shown are 1×10×10 for 96 samples, 1×20×20 for 394 samples, and 1×40×40 for 1536 samples, and when *k* = 20 the grid sizes are 11×3×3 for 96 samples, 24×4×4 for 384 samples, and 96×4×4 for 1,536 samples).

As expected, the General Binary Splitting method consistently required the fewest number of tests in all cases because it is nearly optimal according to group testing theory. Also expectedly, all of the pooling methods were most efficient positive samples were sparse (Fig. 6, top row where *k* = 1). The two non-adaptive methods (DNA Sudoku and 2D Pooling) required the highest number of tests when *k* = 1. DNA Sudoku performed slightly worse than 2D pooling (by one test) owing to the fact that the window sizes were co-prime instead of symmetrical like 2D Pooling. At the maximum number of positive samples for our simulations (*k* = 20), Binary Splitting by Halving performed the worst when the positive rate was high and exceeded individual testing for 96 samples (along with many DNA Sudoku simulations). The number of tests required for our Modified 3-Stage approach typically fell somewhere in the middle except for when the number of total samples and positive samples were highest. When *N* = 1, 536 and *k* = 20, the Modified 3-Stage simulations required only slightly fewer tests, on average, than 2D Pooling, which performed the worst (Table 1).

**Table 1.**
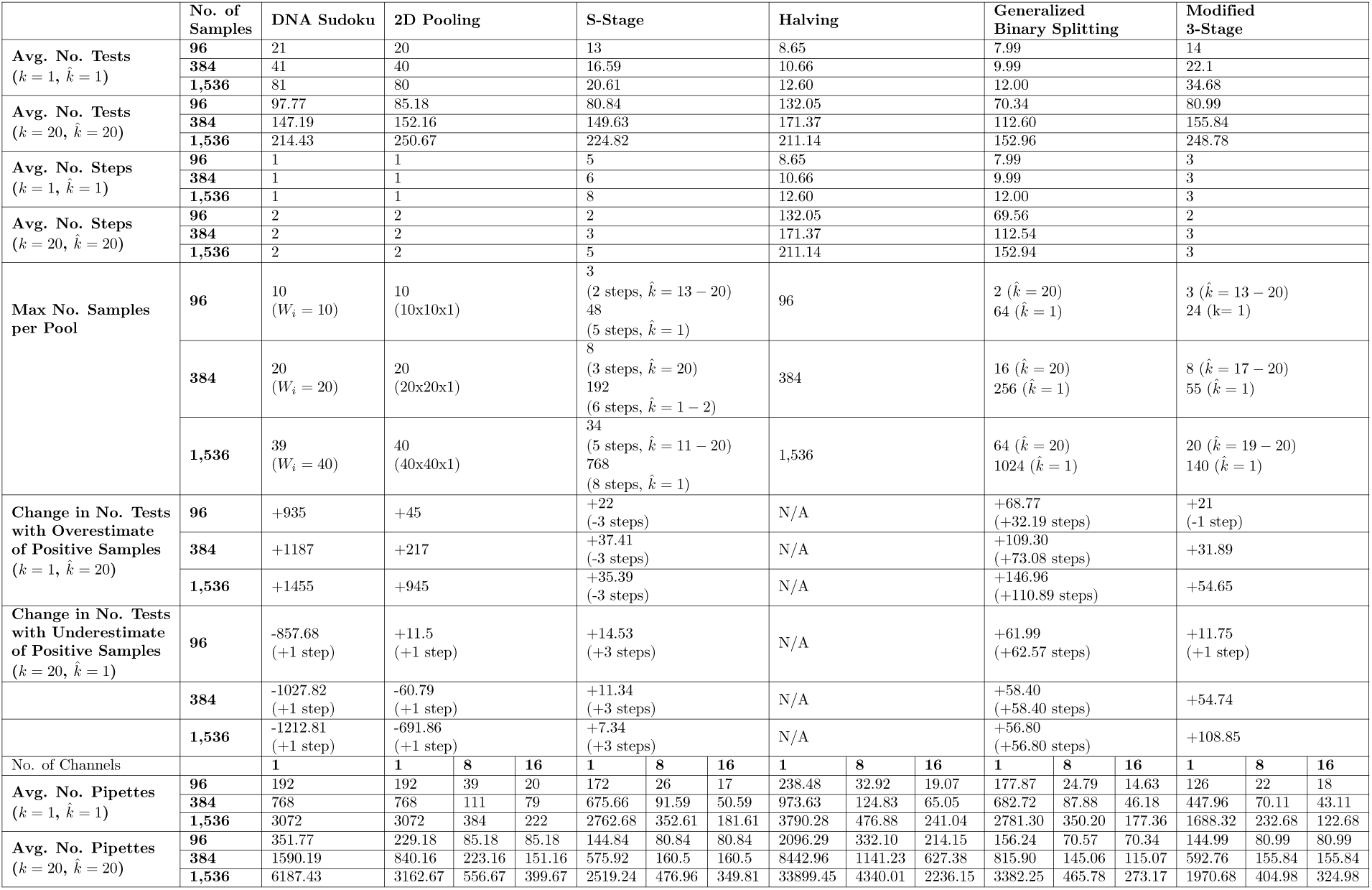
Summary of the performance of pooling methods for each of our areas of interest: number of tests, number of steps, number of samples per pool, robustness, and simplicity. For sample sizes *N* = 96, 384, and 1,536, the table shows the average number for each feature when *k* = 1 and for *k* = 20.

### Number of steps

The number of sequential steps is one of the major factors that differentiates pooling methods. The major benefit of non-adaptive pooling methods is that, in some cases, all of the tests can be run at the same time which means that testing can be completed faster. Clearly, the nonadaptive tests required the fewest number of steps even when the results were ambiguous, necessitating a second round of validation 7. For 96 samples, the highest weight that we tested was 6 which meant that any simulation with 5 or more positive samples was ambiguous. Although higher weights could be used to avoid ambiguous results, the number of tests required would have exceeded individual testing. For 384 and 1,536 samples, up to 9 and 20 positive samples, respectively, were unambiguously identified without exceeding individual testing. The ability to unambiguously identify the positive samples in a single step, however, came with a high cost in the number of tests that needed to be performed. For example, the cost of saving one step when using *w* = 6 versus *w* = 2 for 5 positive samples was on average 56.56 tests for 96 samples. This increased to an additional 1,321.57 tests on average for 1,536 samples using *w* = 21 (1 step) versus *w* = 4 (2 steps) when *k* = 20.

For 2D Pooling, ambiguous results occurred more frequently as the number of positive samples increased and were more likely in larger grid arrangements (Fig. 11). Unlike the DNA Sudoku results, which were always ambiguous or unambiguous based on the weight and the number of positives, ambiguous results for 2D Pooling depended on the random arrangement of the positive samples in the grid and therefore were not always consistent for a given window size. For 96 samples, up to 16 positive samples could be identified in a single step but this only occurred in 1% of the simulations. At 5 positive samples (the highest number that could be unambiguously identified using DNA Sudoku), 66% of the simulations required only one step using 3×3×11 grids (66 tests). For 384 samples, up to 20 positive samples were unambiguously identified using 3×3×43 grids (258 tests) in 15% of the simulations. For 1,536 samples, 20 positive samples were unambiguously identified in 60% of the 3×3×171 grid simulations (1,026 tests), and in only 6% of the 6×6×43 grid simulations (516 tests). This shows that reducing the grid size increases the chances of an unambiguous result but, again, it comes with a large increase in the number of tests (1,026 tests for 3×3×171 grid vs. 160 tests for 20×20×4 at 1,536 samples).

**Fig 7.**
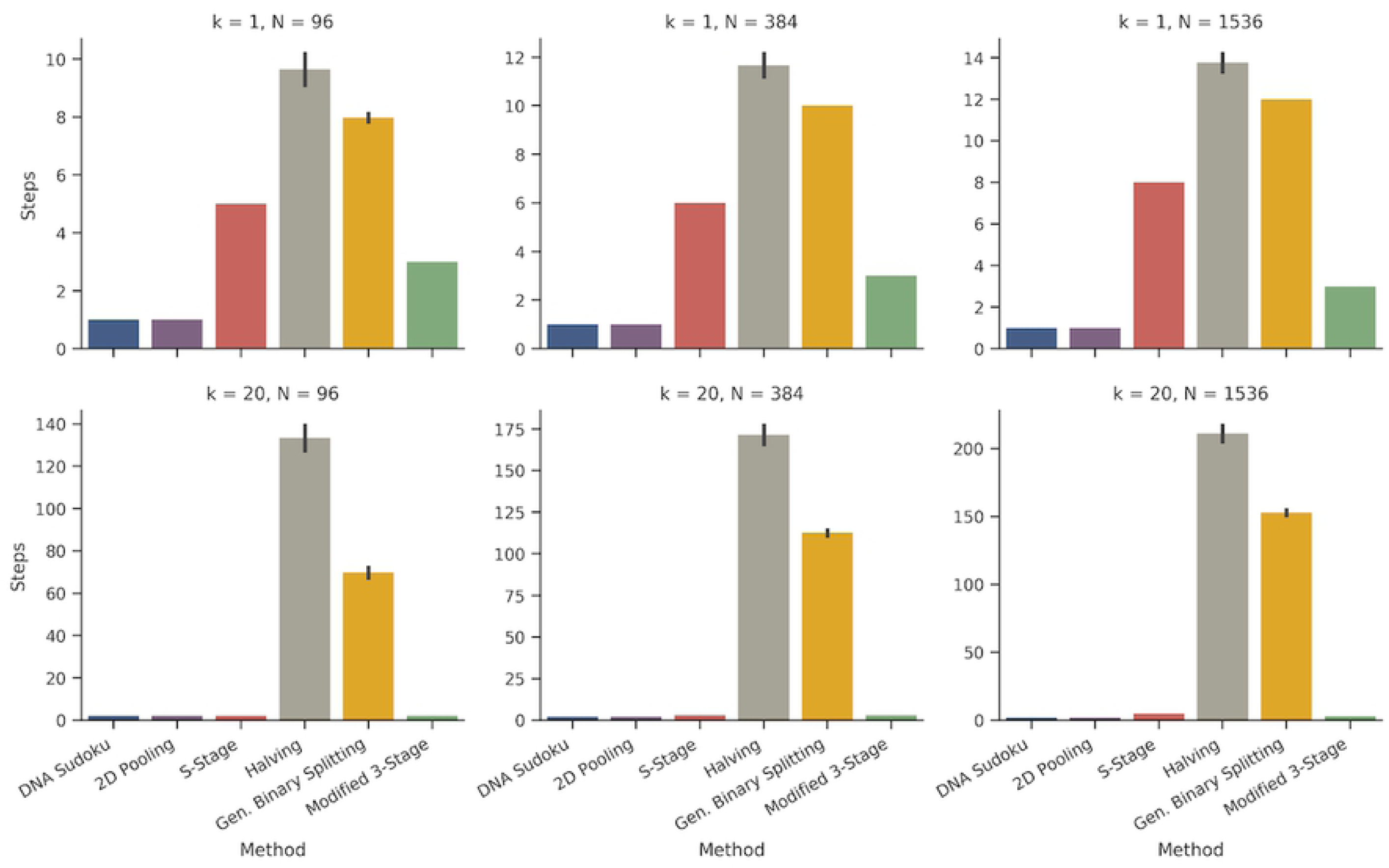
Comparison of the number of steps required for each pooling method. The bar plot shows the average number of steps required for each method and the error bars are the standard deviation across 100 simulations (note the different scales for the y-axis). The top row shows simulations with one positive sample and the bottom row shows simulations with 20 positive samples. The columns are different sample sizes from left to right: 96, 384, and 1,596. These results are from the same set of simulations as shown in Fig 6.

**Fig 8.**
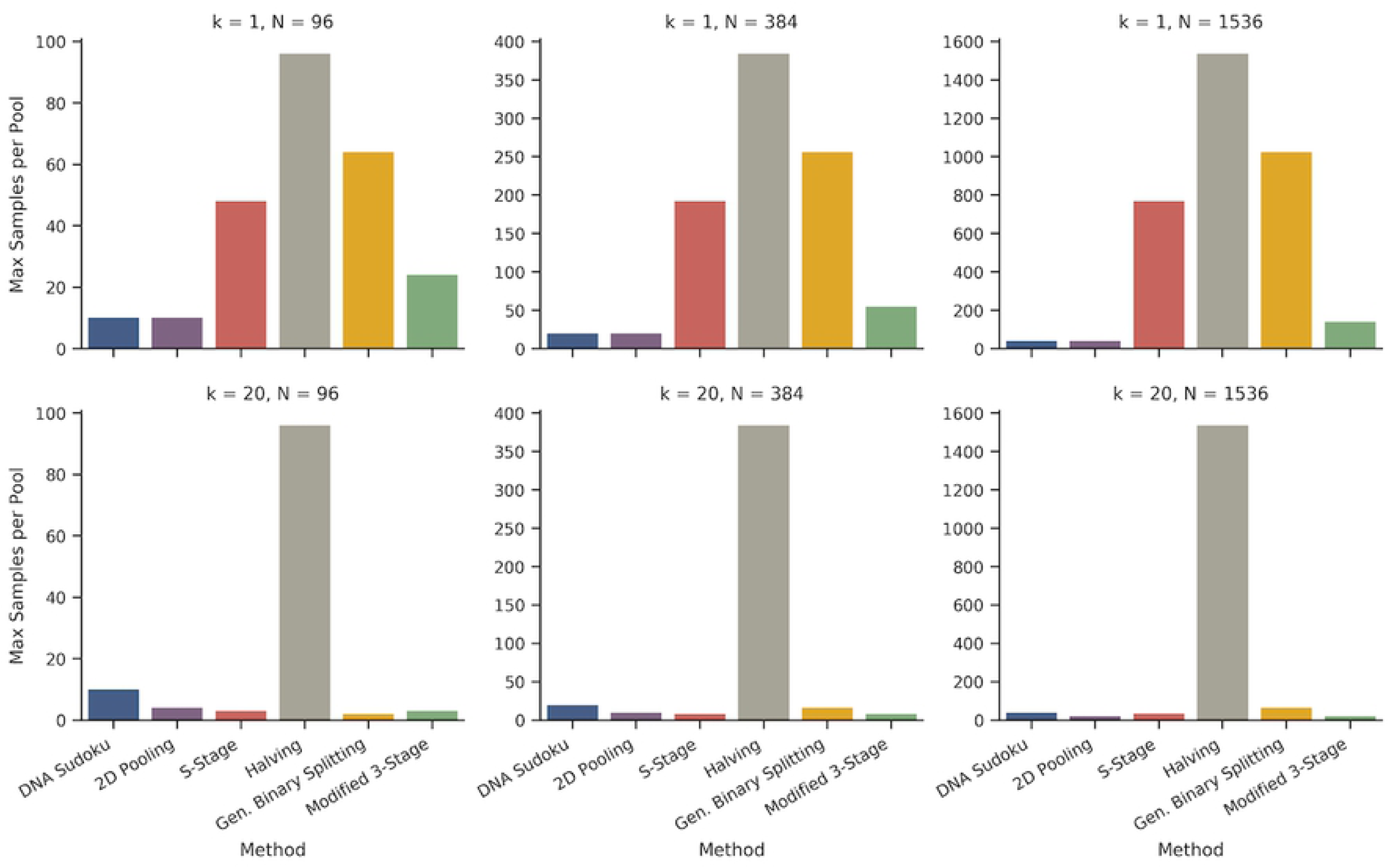
Comparison of the maximum number of samples per pool for each pooling method. The bar graphs plot the maximum number of samples in a single pool for each method (note the different scales for the y-axis). The top row shows simulations with one positive sample and the bottom row shows simulations with 20 positive samples. The columns are different sample sizes from left to right: 96, 384, and 1,596. These results are from the same set of simulations as shown in Fig 6.

**Fig 9.**
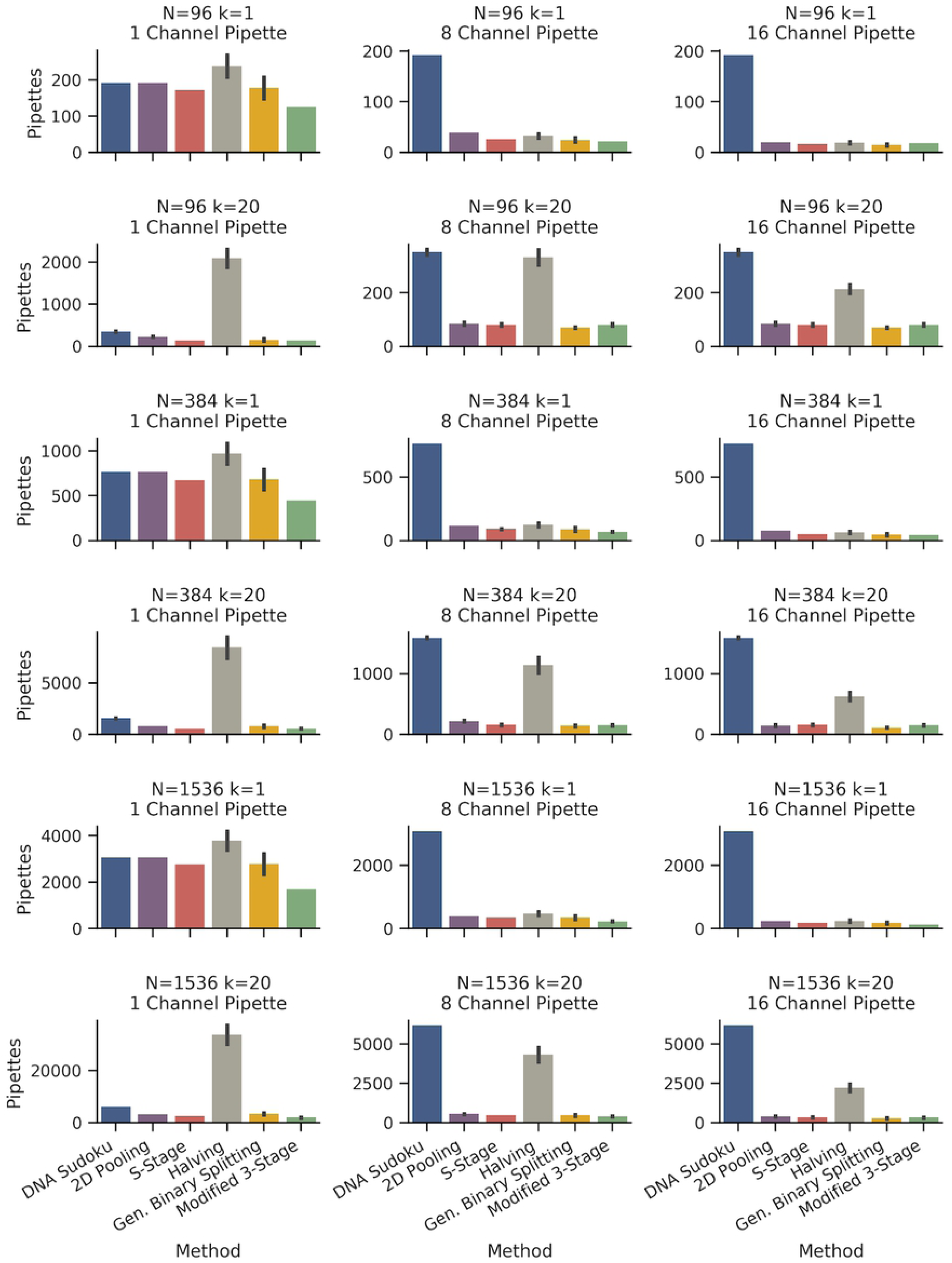
Comparison of the average number of pipettings for each pooling method. The number of pipettings required for each pooling method is an indicator of method simplicity and reproducibility. The bar charts indicate the average number of pipettings required to create pools for each method using 1-, 8-, and 16-channel pipettes (columns). The rows are the results from simulations using N = 96, 384, and 1,536 samples and *k* = 1 or 20 positive samples (note the different scale on the y-axes). The error bars represent the standard deviation of 100 simulations. The DNA Sudoku method does not benefit from the use of multichannel pipettes so the number of pipettings is the same across each row. These results are from the same set of simulations as shown in Fig 6.

**Fig 10.**
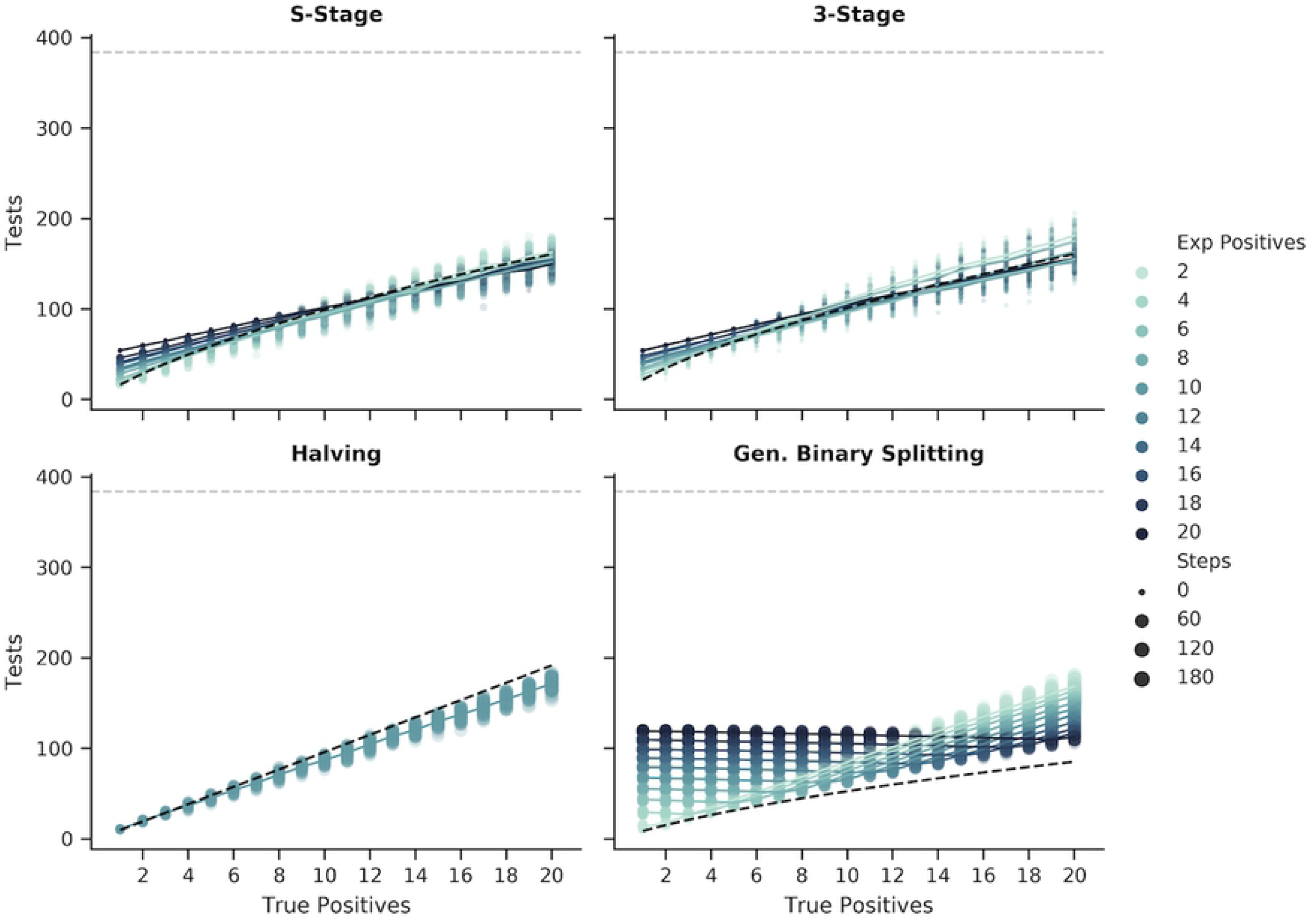
Changes in the number of tests and steps given different estimated positive rates for the adaptive methods. The figure shows the number of tests (y-axis) and the number of steps (marker size) required to recover all positive samples (x-axis) in simulations with *N* = 384 samples using each of the adaptive methods. For each method, except for Binary Splitting by Halving, the pooling scheme was optimized around the expected number of positive samples (marker color) provided to each simulation. Each point represents a single simulation and the lines are the average number of tests for a given number of expected positives. The black dashed line in the S-Stage, 3-Stage, and Binary Splitting by Halving figures represents the upper bound of the number of tests (assuming that the number of positive samples is estimated correctly, where applicable). For the Generalized Binary Splitting figure, the number of tests approaches the lower bound (black dashed line) when 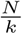 is large.

**Fig 11.**
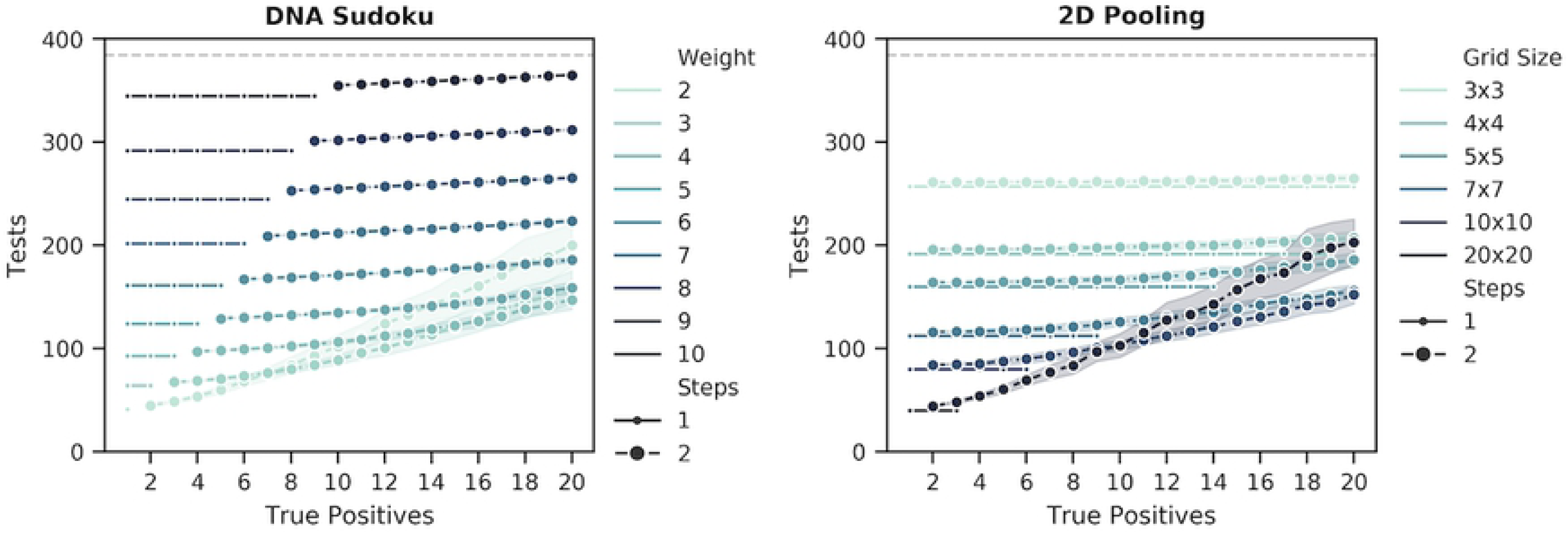
Changes in the number of tests and steps given different estimated positive rates for the non-adaptive methods. The figure shows the number of tests (y-axis) and the number of steps (marker size) required to recover all positive samples (x-axis) in simulations with *N* = 384 samples using DNA Sudoku and 2D Pooling methods. Each point is the average number of tests required for 100 simulations and the width of the bands is the standard deviation. The simulations were run using different weights for DNA Sudoku and different symmetrical 2D grid sizes for 2D Pooling. Small markers indicate unambiguous results that required only a single round of testing and the larger markers indicate ambiguous results that required a second validation step to correctly identify the positive samples. The grey dashed line is the number of tests required for individual testing.

For the adaptive methods, the trade off for being more efficient in the number of tests is often an increase in the number of sequential steps. This is most striking in the case of the Generalized Binary Splitting method which performed the best overall in the number of tests but, in some cases, required over 100 steps. Even so, the Binary Splitting by Halving method required even more steps than the General Binary Splitting method. This is partially due to the fact that the number of tests is highly correlated with the number of steps for both of these methods and the General Binary Splitting algorithm does a better job of minimizing the number of tests (the General Binary Splitting method also switches to individual testing in some cases and, because individual tests can be completed in parallel, this can also reduce the total number of steps). The number of steps required for the S-Stage approach were much more moderate compared to the binary splitting algorithms; however, they did get as high as 8 sequential steps. Our Modified 3-stage approach performed the best among the adaptive methods because it enforced a maximum of 3 steps.

### Number of samples per pool

The number of samples that are combined in a single pool is a very important practical concern because it can determine whether the assay can produce accurate results. Typically, assays can fail to identify positive samples if the positive signal is diluted beyond the limit of detection. This means that pooling approaches that limit the number of samples per pool are more likely to perform better in practice. Fig 8 compares the maximum number of samples per pool for each pooling method.

For the DNA Sudoku simulations, the number of samples per pool was determined by the pooling interval. Because the size of the pooling interval determined the number of pools that the samples would be divided into, a smaller interval resulted in more samples per pool. For 96 samples, the smallest interval was 10 which resulted in up to 10 samples per pool, for 394 samples, the smallest interval was 20 with 20 samples per pool, and for 1,536 samples the smallest interval was 40 with 39 samples per pool (Table 1). For 2D Pooling, the number of samples per pool was equal to the number of samples in each column or row for a given grid layout. The largest grid layouts had the largest number of samples per pool: 10 for 96 samples, 20 for 384 samples, and 40 for 1,536 samples. The number of samples per pool was very consistent for both DNA Sudoku and 2D pooling. For the adaptive approaches, the number of samples per pool varied at each step and tended to be the largest for the first step. For the S-Stage, the Modified 3-stage, General Binary Splitting approaches, the maximum number of samples per pool was lowest when the number of expected positive samples was high and increased as the number of expected positive samples decreased (Table 1). The Modified 3-Stage approach always had the same or fewer samples per pool compared to the S-stage approach. This was particularly true when the number of positive samples was low, resulting in a reduction of 24, 137, and 628 samples per pool for N = 96, 384, 1,536, respectively. The Binary Splitting by Halving method required the largest number of samples per pool at 96, 384, and 1536, due to the need to pool and test all of the samples together as the first step.

### Simplicity of pooling method

The simplicity of a pooling method can be somewhat subjective. However, one of the major points of failure when combining pools by hand, is mistakes in pipetting the wrong samples. Therefore, we used the number of individual pipetting actions required for each method using 1-, 8-, and 16-channel pipettes as an indicator for the simplicity and reproducibility of each method. Fig 9 compares the number of pipettings required for each method with either 1 or 20 positive samples for each of the sample sizes.

In most methods, using a multichannel pipette reduced the number of pipettings by an order of magnitude in some cases. Compared to the 8-channel pipette, the channel pipette reduced the number of pipettings only for schemes where the size of the pools were large. Our Modified 3-Stage method required the fewest pipettings with a single-channel pipette, compared to the other methods; however, the S-Stage method performed similarly well in cases where the number of steps happened to be similar (i.e. in Fig 9, both methods required the same number of steps when *k* = 20 for *N* = 96 and 384). This generally translated into the best performance when using multichannel pipettes; although, the General Binary Splitting simulations performed slightly better in cases where the size of the pools were large, making the multichannel pipettes more efficient.

Binary Splitting by Halving is the least efficient method in number of pipettings, likely because the method requires many samples to be pooled at each step for many steps. The performance is slightly improved when multichannel pipettes are used but it is still the least efficient in many cases. Using a single pipette, DNA Sudoku was not the most inefficient compared to the other methods. However, because the samples that are combined in each pool are spaced out in different intervals instead of in consecutive groups, the number of pipettings did not improve by using multichannel pipettes. This means that, in the best case, a laboratory technician would need to correctly pipette ∼ 200 (*N* = 96) to ∼ 6, 000 (*N* = 1, 536) times to combine the samples into pools.

### Sensitivity to misestimation of positive rate

Most of the methods described here required an estimate of the positive rate in order to design the pooling scheme. In some cases, over or underestimating the number of positives can have large impacts on the number of tests and/or the number of steps required to complete the assay. Methods that minimize the impact of misestimation are more robust to fluctuating rates in the sample population which is common in outbreak scenarios.

Binary Splitting by Halving was, by default, the most robust of the approaches because the protocol was not modified based on any estimate of the number of positive samples (Fig 10, second from right). Although the number of tests increased when there were more positive samples, assuming a fixed sample size, knowledge of the number of positive samples did not have any impact on the performance of this approach. In contrast, the size of the initial pool for the Generalized Binary Splitting method depended on 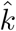 and, among the adaptive approaches, misestimation of the true value resulted in the largest impact on the number of tests and steps (Fig 10, right). The consequences of extreme overestimation (*k* = 1 and 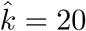) and underestimation (*k* = 20 and 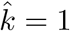) are provided in Table 1 which shows that method is more sensitive to overestimation than to underestimation. Of the methods that depended on an estimate of the number of positive samples, the S-Stage (Fig 10, left) and our Modified 3-Stage approach (Fig 10, second from left) were the most robust to misestimations of *k*. The number of steps was more robust in the Modified 3-Stage approach than the S-Stage due to the 3-Step constraint; however, the Modified 3-Stage was more sensitive in the number of tests in some cases (Table 1).

DNA Sudoku (Fig 11, left) was the most sensitive method overall. Overestimating of the number of positive samples caused the weight of the pooling design 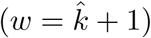 to be set higher than it needed to be. When this happened, all of the positive samples were still unambiguously identified but each unnecessary increase in the weight required more than 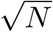 additional tests. When the number of positive samples was underestimated, fewer tests were performed but the pooling scheme was no longer able to unambiguously identify the positive samples in a single step and a second round of verification was required. A similar pattern occurred in the 2D Pooling simulations (Fig 11, right). While the grid dimensions did not directly depend on *k*, generally larger grids were more efficient when the number of positive samples was low and smaller grids reduced ambiguous results when the number of positive samples was high but at the cost of many more tests. However, because 2D Pooling was constrained to two dimensions, the number of tests did not vary as drastically as DNA Sudoku.

### Experimental validation of modified 3-Stage approach

To experimentally validate our modified 3-Stage approach, we set up a controlled experiment with *C. burnetti* DNA spiked into complex microbiome background samples. All of the 24 background samples used in the validation experiment were negative for the *C. burnetti* pathogen prior to the spike-in and each background extraction was found to have similar amounts of the 16S gene (CT values of 29 to 31), indicating similar background bacterial loads. *C. burnetti* was detected in the spike-in samples prior to pooling. The random placement of the *C. burnetti* positive samples within the eight 96-well plates is shown in Table 2. Although the expected number of positive samples per plate was ∼2 given the 1.3% carriage rate, the actual number of positives ranged from 0 to 3 and none of the plates had exactly 2 positive samples (Table 2). The TaqMan assay was able to accurately identify the positive pools without any false positives or false negatives even during the first step when the number of samples per pool was the largest at 16. Using an 8-channel pipette where appropriate, a total of 180 pipettings was required to pool the samples. A total of 120 TaqMan assays were performed which is ∼84% fewer than would be required to individually test 768 samples.

**Table 2.** Using the modified 3-Stage approach we were able to accurately recover all of the positive *C. burnetii* samples. Eight 96-well plates were filled with background DNA from complex cow milk microbiome samples and 10 randomly chosen samples had *C. burnetii* DNA spiked in. The table shows the number and location of the positive samples in each 96-well plate and the number of tests and pipettings required to identify the positive samples using our Modified 3-Stage pooling approach.

| Plate # | Positive Samples (k) | Exp. Positive Samples ($\hat{k}$) | Samples (N) | Positive Wells | Tests | Pipettings |
| --- | --- | --- | --- | --- | --- | --- |
| 1 | 3 | 2 | 96 | F1, A4, G9 | 30 | 48 |
| 2 | 1 | 2 | 96 | H2 | 14 | 20 |
| 3 | 1 | 2 | 96 | D9 | 14 | 20 |
| 4 | 1 | 2 | 96 | E11 | 14 | 20 |
| 5 | 0 | 2 | 96 | NA | 6 | 12 |
| 6 | 3 | 2 | 96 | F3, A10, C10 | 22 | 28 |
| 7 | 0 | 2 | 96 | NA | 6 | 12 |
| 8 | 1 | 2 | 96 | D4 | 14 | 20 |

## Conclusions

Picking the right pooling approach for a given pathogen surveillance campaign can be a complicated decision, which is often driven by a set of conflicting constraints and priorities, including budgetary limitations, complexity of the procedure (and thus likelihood of human error), and time-to-answer requirements. As is evident from the data presented in this manuscript, no single group theory approach is a clear winner under all these possible constraints – the correct choice depends on the predominant constraints placed on the surveillance campaign. Below we present some of the practical implications of the various group theory approaches outlined in this manuscript.

When minimizing the total number of tests is the absolute overriding goal and time-to-answer is not an important constraint, Generalized Binary Splitting is the optimal choice. This approach minimized the total number of tests while maintaining a reasonable complexity, as measured by the number of distinct pipetting actions, but sacrifices speed due to significantly increase in the number of serial steps. On the other hand, when speed is the predominant constraint, DNA Sudoku can offer a single-step pooling approach with the minimum number of tests, at the cost of significant complexity. DNA Sudoku, however, is far from optimal for monitoring rapidly changing pandemics due to its extreme sensitivity to misestimation of the carriage rate of the pathogen in population.

A good middle ground between the adaptive and non-adaptive pooling approaches is the Modified 3-Stage approach – our preference in our own surveillance applications. While it is never the absolute best in any one category, it is always nearly optimal in terms of number of serial steps (2nd best), complexity (2nd best), number of tests (4th best), and extremely resilient to misestimation of the carriage rate (2nd best). The latter is particularly important, as it allows this approach to be useful for surveillance in situations with rapidly changing pathogen carriage rates (e.g. in pandemic or seasonal outbreaks), while keeping number of serial steps as low as possible for an adaptive method.

